# AAV-Mediated Dual-Gene Therapy Restores Metabolic Function in Mice with Propionic Acidemia

**DOI:** 10.64898/2026.03.06.709717

**Authors:** Huixia Xu, Zewen Tao, Ting Zhang, Xuli Zhang, Ying Zhou, Ziyan Cen, Jie Liu, Hao Zhang, Alimujiang Maimaitijiang, Dan Chen, Dali Li, Shuming Yin, Lei An, Xinwen Huang, Yue Zhang

**Author notes:** **Corresponding authors:** Lei An, No.115, Ximen Avenue, Huaihe Hospital of Henan University, Kaifeng 475000, Henan, P. R. China., Xinwen Huang, The Children’s Hospital, Zhejiang University School of Medicine, Liangzhu Laboratory, Hangzhou 310000, China., Yue Zhang, The Children’s Hospital, National Clinical Research Center for Child Health, Zhejiang University School of Medicine, Liangzhu Laboratory, Hangzhou 310000, China., Shuming Yin, Shanghai Frontiers Science Center of Genome Editing and Cell Therapy, Shanghai Key Laboratory of Regulatory Biology, School of Life Sciences, East China Normal University, Shanghai 200241, China. These authors contributed equally to this study (Huixia Xu, Zewen Tao, Ting Zhang). These authors share senior authorship (Shuming Yin, Lei An, Xinwen Huang, Yue Zhang).

## Abstract

**Background and Aims:** Propionic acidemia (PA) is a rare autosomal recessive disorder caused by mutations in *PCCA* or *PCCB*, which encode the two subunits of propionyl-CoA carboxylase (PCC). PCC deficiency causes toxic metabolite accumulation and multi-organ damage. Current management, including dietary restriction, pharmacological support, and liver transplantation, does not restore enzymatic activity. We developed a dual-gene adeno-associated virus (AAV) therapy that delivers both PCC subunits to treat both PA subtypes.

**Methods:** We generated a clinically relevant *PCCA*-R73W knock-in mouse model and administered AAV8 vectors encoding native human *PCCA* and *PCCB* under the control of a liver-specific thyroxine-binding globulin promoter (AAV8-TBG-h*PCCA*-P2A-h*PCCB*). Metabolite levels and organ safety were longitudinally assessed.

**Results:** Dual-gene therapy produced dose-dependent reductions in plasma C3/C2 ratio, 3-hydroxypropionic acid, 2-methylcitric acid, and propionylglycine, and significantly outperformed single-gene (*PCCA*-only) therapy. Neonatal facial-vein injection achieved metabolic correction comparable to or better than adult treatment. The longitudinal follow-up revealed sustained efficacy over a 16-week period, with no signs of hepatotoxicity or adverse effects.

**Conclusions:** Single-dose, dual-gene AAV therapy achieves sustained metabolic correction and demonstrates long-term safety in a clinically relevant PA model, supporting its translational potential for both type I and type II propionic acidemia.

## Introduction

Propionic acidemia (PA) is an autosomal recessive inborn error of metabolism caused by pathogenic variants in *PCCA* or *PCCB*, which encode the α and β subunits of the mitochondrial enzyme propionyl-CoA carboxylase (PCC)[1–3]. The loss of PCC activity results in the accumulation of propionylcarnitine (C3), 3-hydroxypropionic acid (3-HP), and 2-methylcitric acid (2-MeCit), driving recurrent metabolic acidosis, hyperammonemia, neurological injury, and multi-organ dysfunction[4,5]. Most patients present during the neonatal period or early infancy, and the risk of acute metabolic decompensation persists throughout life[6,7].

PCC is a heterododecameric enzyme (∼750 kDa) composed of six alpha (*PCCA*) and six beta (*PCCB*) subunits that function as an obligate complex[1,3]. *PCCB* is unstable in the absence of *PCCA*; as a result, patients *with PCCA* deficiency (type I PA) typically lack both subunits[8,9]. Patients with PCCB deficiency (type II PA) can similarly show reduced *PCCA* protein levels [1,2,10]. Mechanistically, heterooligomeric assembly into the dodecamer appears to be critical for stabilizing each subunit; without the partner subunit, the non-associated protein is prone to degradation. This mutual dependency means that replacing only one subunit may not restore balanced stoichiometry or full enzymatic activity, thus limiting single-gene strategies for both PA subtypes.

Recent nucleic acid therapeutics have broadened the treatment options for PA. Jiang et al. showed that dual mRNA therapy encoding both PCC subunits restored hepatic PCC enzyme activity and lowered circulating biomarkers, including 2-MeCit and 3-HP, in a PA mouse model over 3- and 6-month dosing periods[11]. However, plasma biomarkers did not fully normalize to wild-type levels, and the transient nature of mRNA requires repeated dosing to maintain benefit, representing a significant practical hurdle for a chronic pediatric condition[12,13].

Adeno-associated virus (AAV) vectors are well suited for the treatment of genetic diseases, given their safety record, low immunogenicity, and ability to sustain transgene expression after a single dose[14,15]. In this study, we packaged both CDS of human *PCCA* and *PCCB* into a single AAV vector and tested this dual-gene therapeutic strategy in a clinically relevant knock-in mouse model. We demonstrated that the synergistic delivery of dual-gene therapy results in superior therapeutic outcomes compared to single-gene therapy, maintaining its efficacy over the long term without any significant side effects. Neonatal treatment in mice yields more significant therapeutic advantages than when administered at 4 weeks of age.

## Results

The *PCCA* c.229C>T (p.R77W) variant is recurrently identified in patients with PA and has been reported as a pathogenic mutation associated with the classic form of the disease[8,16,17]; however, its functional consequences have not been characterized in an animal model. To fill this gap, we introduced the corresponding murine *Pcca* R73W substitution via CRISPR/Cas9-mediated knock-in (Fig. S1A, B) and characterized the resulting *PCCA^KI/KI^* mice. Compared with WT and heterozygous littermates, homozygous animals showed clear growth retardation, with reduced body size and persistently lower body weight throughout postnatal development (Fig. 1A, B). TEM of liver tissue revealed marked mitochondrial abnormalities in *PCCA^KI/KI^* mice, including swelling and loss of cristae organization (Fig. 1C). In addition to growth impairment and canonical biomarker elevations, *PCCA^KI/KI^* mice exhibited hyperammonemia, elevated propionylglycine and amino acids, and mitochondrial ultrastructural injury in cardiomyocytes (Fig. 1C). The *PCCA^KI/KI^* model thus recapitulates the key features of human PA: impaired growth, hepatic mitochondrial damage, and sustained metabolic derangement. Beyond growth impairment and the elevation of canonical biomarkers, *PCCA^KI/KI^* mice also showed hyperammonemia, increased levels of propionylglycine and amino acids, and mitochondrial ultrastructural damage in cardiomyocytes (Fig. S1C-H).

**Figure 1.**
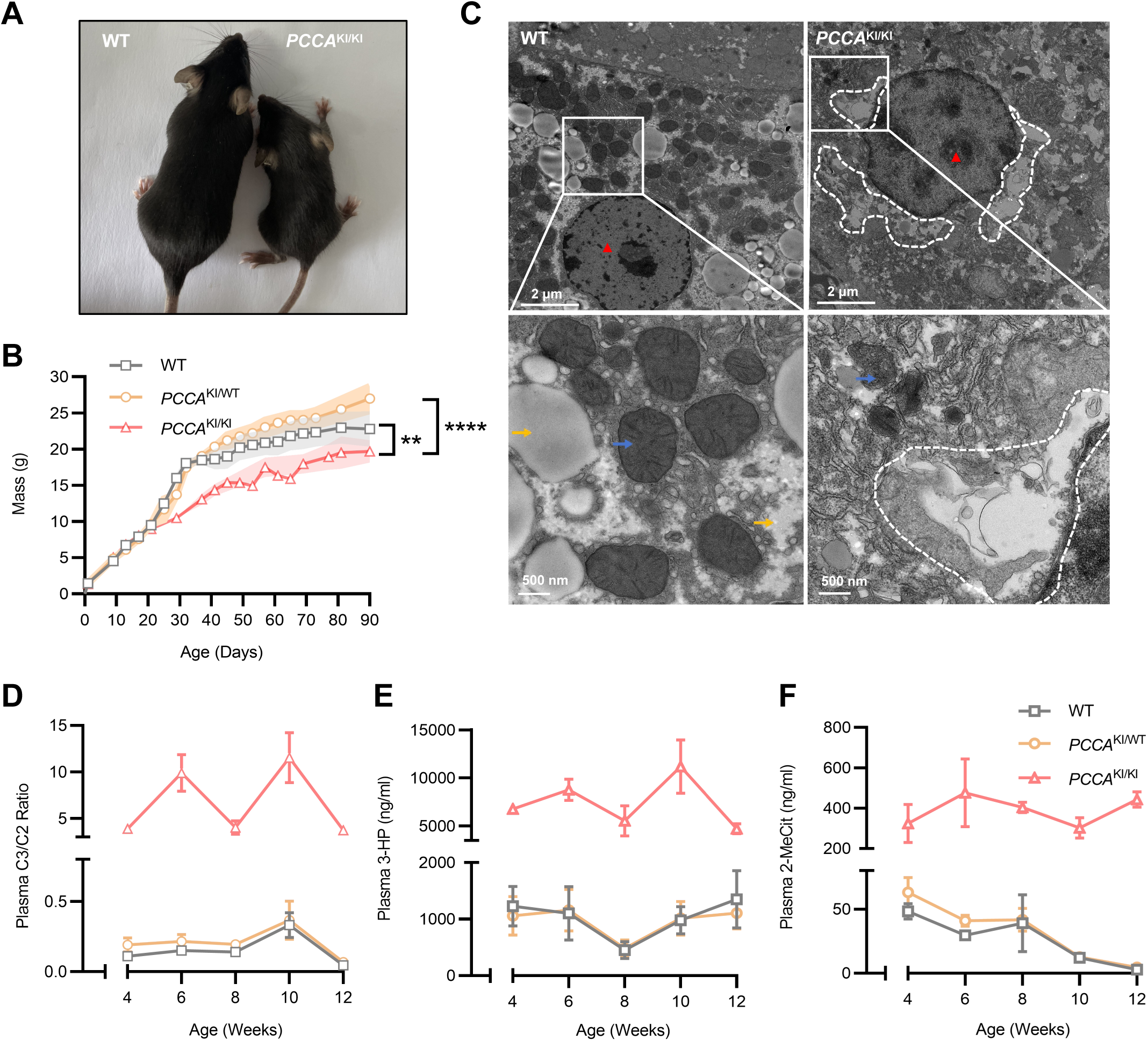
*PCCA* knock-in mice exhibit growth retardation, mitochondrial ultrastructural injury in the liver, and elevated circulating PA biomarkers. (A) Representative gross appearance of age-matched wild-type (WT) and homozygous *PCCA* knockout (*PCCA*^KI/KI^) mice, illustrating reduced body size in *PCCA*^KI/KI^ animals. (B) Longitudinal body weight curves of WT, heterozygous (*PCCA*^KI/WT^), and homozygous (*PCCA*^KI/KI^) mice from birth to postnatal day 90, demonstrating impaired growth in *PCCA*^KI/KI^ mice. (C) Transmission electron microscopy (TEM) of liver tissue (top and bottom rows, lower and higher magnification, respectively) showing abnormal mitochondrial morphology in *PCCA*^KI/KI^ mice compared with WT, including prominent mitochondrial swelling and disrupted cristae architecture (dashed outline), with representative regions highlighted. (D-F) Ultra-Performance Liquid Chromatography - Mass Spectrometry / Mass Spectrometry (UPLC-MS/MS) analysis of every two-week blood samples revealed sustained elevations of canonical propionylcarnitine/acetylcarnitine (C3/C2) (D), 3-hydroxypropionic acid (3-HP) (E), and 2-methylcitric acid (2-MeCit) (F) biomarkers in *PCCA^KI/KI^* mice, whereas heterozygous mice exhibited levels comparable to WT. 3-HP and 2-MeCit are reported in ng/mL, and C3/C2 is shown as a ratio. Data points represent group averages ± SEM. \**p* < 0.05; \*\**p* < 0.01; \*\*\**p* < 0.001; \*\*\*\**p* < 0.0001 (one-way ANOVA followed by the Tukey’s posttest). Sample sizes are indicated in the corresponding panels.

Next, we asked whether the dual-gene construct produced functional proteins that reached mitochondria. MEFs isolated from *PCCA^KO/KO^* embryos were transduced with a lentiviral vector encoding codon-optimized human *PCCA* and *PCCB* linked by a P2A peptide, with sfGFP as a reporter (Fig. 2A); vimentin staining confirmed the mesenchymal identity of all MEF lines used (Fig. S2A). RT-qPCR demonstrated marked upregulation of both *PCCA* and *PCCB* mRNA in transduced *PCCA^KO/KO^* MEFs compared with untransduced controls (Fig. S2B,C). At the protein level, confocal imaging showed strong expression of both PCCA and PCCB in transduced cells, at levels comparable to endogenous PCC in WT MEFs (Fig.2B). The dual-gene cassette thus directs the expression and mitochondrial targeting of both PCC subunits *in vitro.* In Lenti-sfGFP-transduced *PCCA^KO/KO^* cells, PCCA was undetectable and PCCB signal was reduced relative to WT, consistent with the known instability of PCCB without its PCCA partner [8,9]. The dual-gene cassette thus directs expression and mitochondrial targeting of both PCC subunits *in vitro*.

**Figure 2.**
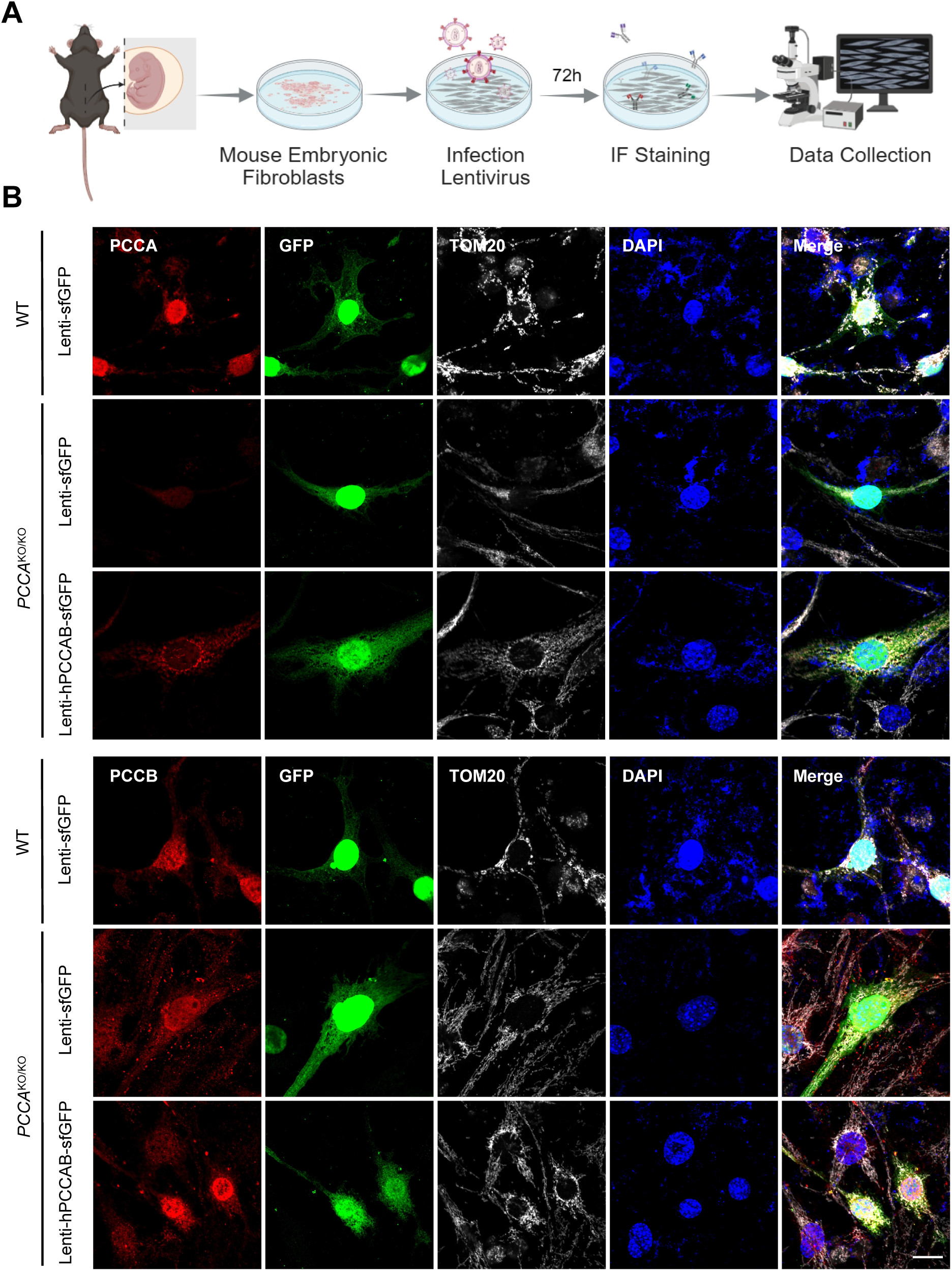
Localization of lentivirus-delivered human PCCA and PCCB proteins to mitochondria in PCCA-deficient mEFs. (A) Schematic of the experimental workflow: Mouse embryonic fibroblasts (MEFs) isolated from *PCCA*^KO/KO^ embryos were transduced with lentivirus encoding human *PCCA* and *PCCB* (Lenti-h*PCCA*/B) or SFGFP alone (Lenti-SFGFP), followed by immunofluorescence staining and confocal imaging (Olympus FV3000). (B) Representative confocal images of wild-type (WT), Lenti-SFGFP-transduced, and Lenti-h*PCCA*/B-transduced MEFs immunostained for *PCCA* (top panels, red) or *PCCB* (bottom panels, red), together with SFGFP (green), the mitochondrial marker TOM20 (white), and DAPI (blue). Merged images demonstrate the co-localization of exogenous *PCCA* and *PCCB* with TOM20 in Lenti-h*PCCA*/B-transduced cells, confirming the correct mitochondrial targeting of the transgene-derived proteins. Scale bar, 5μm.

To achieve better targeting, we selected a combination of the AAV8 serotype and human thyroxine-binding globulin promoter (TBG)[18,19]. We initially compared the *in vivo* performance of codon-optimized versus native (non-codon-optimized) human PCCA/PCCB sequences. AAV8 vectors with native sequences resulted in significantly greater reductions in both the plasma C3/C2 ratio and 3-HP compared to their codon-optimized counterparts (Fig. S3A, B). Consequently, native sequences were employed in all subsequent experiments.

PCCA^KI/KI^ mice were administered a single intravenous injection of AAV8-TBG-hPCCA-hPCCB (native sequences) at three different dose levels (1×10¹¹, 5×10¹¹, and 1×10¹² vg/mouse) when they were four weeks old. Plasma metabolites were measured four weeks later using Ultra-Performance Liquid Chromatography - Mass Spectrometry / Mass Spectrometry (UPLC-MS/MS) (Fig. 3A). At both time points, the mid- and high-dose groups exhibited significantly greater reductions in PA biomarkers compared to the low-dose group across nearly all analytes tested (Fig. 3B-E; Fig. S3C-J). However, the high dose did not consistently outperform the mid-dose; for instance, 3-HP levels were lower in the mid-dose group than in the high-dose group (Fig. 3E). This pattern persisted: at 2 months, the mid- and high-dose groups continued to show significant correction of the C3/C2 ratio, propionylglycine, glycine, and 2-MeCit relative to untreated controls (Fig. S3E, H-J). Weighing the sustained therapeutic benefit against potential dose-related risks, we chose the mid-dose (5×10¹¹ vg/mouse) for all subsequent experiments.

**Figure 3.**
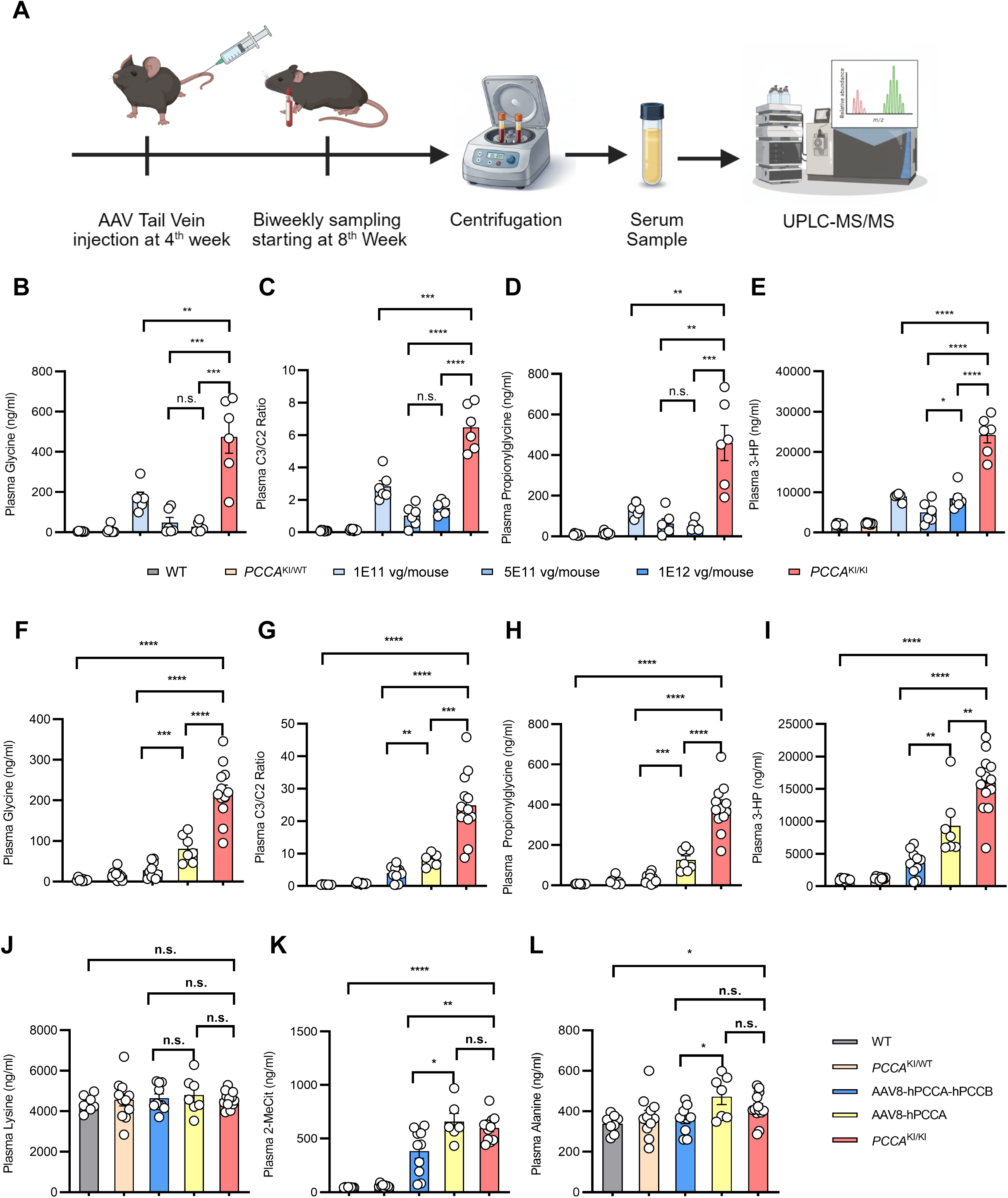
Dual-gene AAV therapy using native human sequences produces a dose-dependent correction of circulating PA biomarkers and outperforms single-gene therapy. (A) Schematic of the in vivo study design: *PCCA*^KI/KI^ mice received a single tail-vein injection of AAV8-TBG-h*PCCA*-h*PCCB* encoding native (non-codon-optimized) human *PCCA* and *PCCB* sequences at 4 weeks of age; plasma was collected at 8 weeks and analyzed by ultra-performance liquid chromatography-tandem mass spectrometry (UPLC-MS/MS). Native sequences were selected based on their superior in vivo performance over codon-optimized variants (see Fig. S3A, B). (B-E) Dose-response analysis of dual-gene AAV therapy (1×10¹¹, 5×10¹¹, and 1×10¹² vg/mouse) on plasma glycine (B), the C3/C2 (propionylcarnitine/acetylcarnitine) ratio (C), propionylglycine (D), and 3-hydroxypropionic acid (3-HP) (E), compared with WT, *PCCA*^KI/WT^, and untreated *PCCA*^KI/KI^ controls. (F-I) Head-to-head comparison of dual-gene (AAV-hPCCA-hPCCB) versus single-gene (AAV-h*PCCA*) therapy at 5×10¹¹ vg/mouse on plasma glycine (F), C3/C2 ratio (G), propionylglycine (H), and 3-HP (I). (J-L) Additional metabolite profiling comparing dual-gene and single-gene therapy for plasma lysine (J), 2-methylcitric acid (2-MeCit) (K), and alanine (L). Data are presented as mean ± SEM. \**p* < 0.05; \*\**p* < 0.01; \*\*\**p* < 0.001; \*\*\*\**p*< 0.0001 (one-way ANOVA followed by Tukey’s posttest). Sample sizes are indicated in the corresponding panels.

In a direct comparison, dual-gene therapy significantly outperformed single-gene (AAV-hPCCA) therapy in reducing glycine, the C3/C2 ratio, and propionylglycine, with the latter achieving only partial correction (Fig. 3F-H). While both approaches decreased 3-HP levels, the reduction was notably more pronounced with dual-gene treatment (Fig. 3I). Thus, co-delivery of both subunits in a single vector offers more comprehensive and profound metabolic correction than PCCA monotherapy, underscoring the necessity of balanced expression of both PCC subunits for complete biochemical rescue. 3K, L). Lysine levels remained consistent across all groups (Fig. 3J). Thus, co-delivering both subunits in a single vector offers a more comprehensive and profound metabolic correction than PCCA monotherapy, underscoring the necessity of balanced expression of both PCC subunits for complete biochemical rescue.

We conducted long-term monitoring of the metabolites in mice injected at 4 weeks of age. In untreated *PCCA*^KI/KI^ mice, circulating glycine, propionylglycine, C3/C2 ratio, 2-MeCit, and 3-HP levels progressively increased with age (Fig. 4A, C, E–G). Treated animals maintained much lower levels of these metabolites throughout the 16-week observation period, remaining significantly below untreated controls at every time point. Alanine and lysine showed more modest separation between groups (Fig. 4B, D).

**Figure 4.**
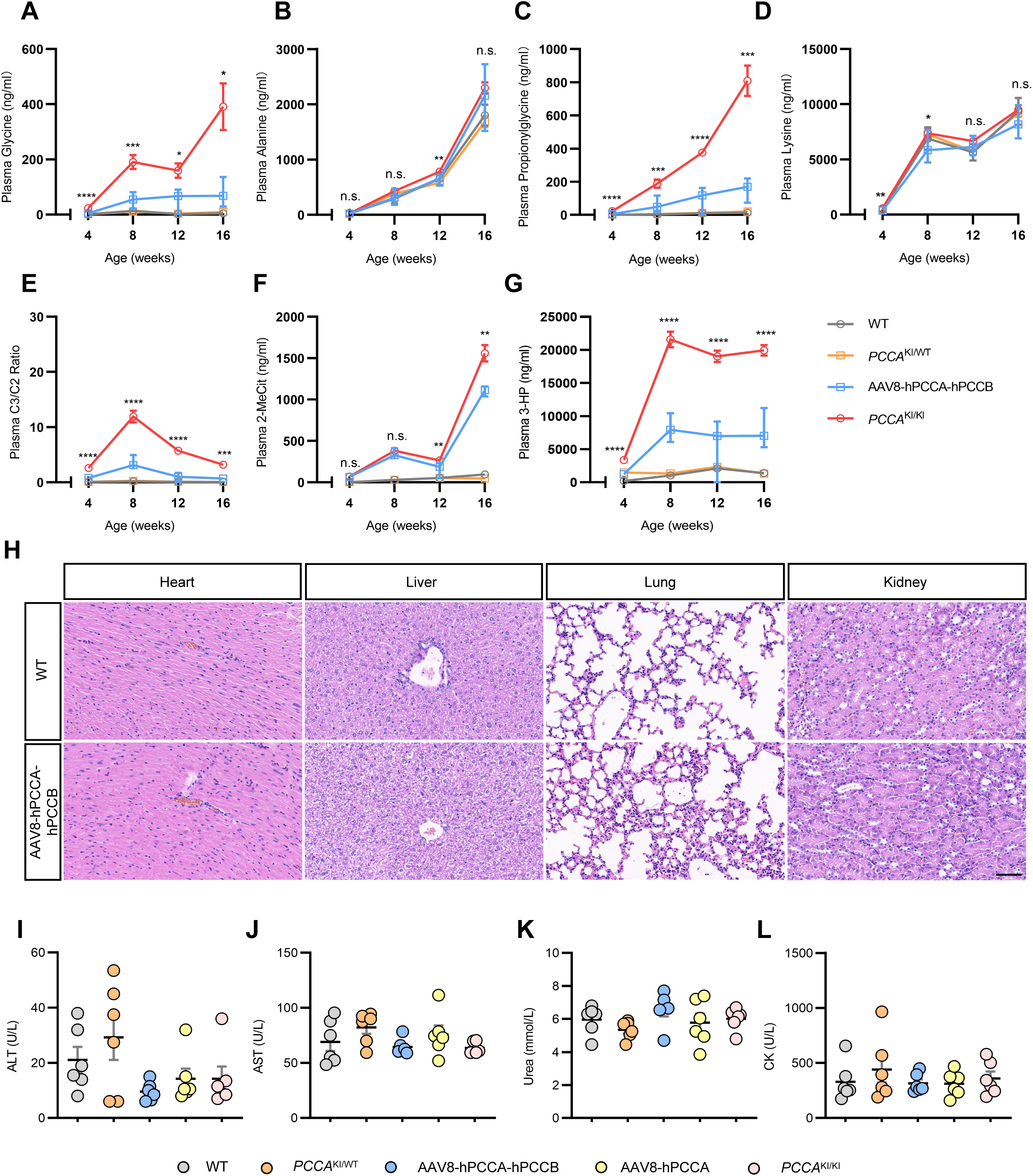
Dual-gene AAV therapy achieves sustained long-term metabolic correction with a favorable safety profile. (A-G) Longitudinal profiling of circulating PA biomarkers from 4 to 16 weeks of age following a single tail-vein injection of AAV8-TBG-hPCCA-hPCCB at 4 weeks: plasma glycine (A), alanine (B), propionylglycine (C), lysine (D), C3/C2 (propionylcarnitine/acetylcarnitine) ratio (E), 2-methylcitric acid (2-MeCit) (F), and 3-hydroxypropionic acid (3-HP) (G) in WT, *PCCA*^KI/WT^, AAV8-h*PCCA*-h*PCCB*-treated, and untreated *PCCA*^KI/KI^ mice. p.i.: post injection. (H) Representative hematoxylin and eosin (H&E)-stained sections of the heart, liver, lung, and kidney from WT and AAV8-hPCCA-hPCCB-treated *PCCA*^KI/KI^ mice at 4 weeks post-injection (8 weeks of age), showing no apparent histopathological abnormalities in treated animals. (I-L) Serum safety biomarkers at 4 weeks post-injection: alanine aminotransferase (ALT, U/L) (I), aspartate aminotransferase (AST, U/L) (J), urea (mmol/L) (K), and creatine kinase (CK, U/L) (L), compared across WT, *PCCA*^KI/WT^, AAV8-hPCCA-hPCCB (dual-gene), AAV8-hPCCA (single-gene), and *PCCA*^KI/KI^ groups. All metabolite concentrations are reported in ng/mL, except for C3/C2(ratio), urea (mmol/L), and enzymatic activities (U/L). Data are presented as the mean ± SEM.\**p* < 0.05; \*\**p* < 0.01; \*\*\**p* < 0.001; \*\*\*\**p* < 0.0001 (two-way ANOVA for A-G; one-way ANOVA followed by Tukey’s posttest for I-L). Sample sizes are indicated in the corresponding panels.

For safety assessment, H&E staining of heart, liver, lung, and kidney tissues at 4 weeks post-injection showed normal tissue architecture in treated *PCCA^KI/KI^* mice, comparable to that in WT (Fig. 4H). Serum ALT, AST, urea, and CK were all within the WT and heterozygous range, with no elevations suggestive of liver, kidney, or muscle toxicity (Fig. 4I-L). Single-gene therapy also showed no safety signals, indicating that the AAV8-TBG platform is well tolerated at these doses. We conducted the biochemical safety panel again at 8 and 12 weeks after injection. At both intervals, all markers in mice treated with either dual-gene or single-gene therapy remained comparable to those in WT and heterozygous controls, showing no signs of delayed toxicity (Fig. S4A-H). A single dose of dual-gene AAV at 4 weeks of age thus provides at least 4 months of metabolic correction without detectable organ toxicity.

We then asked whether earlier intervention would improve metabolic correction.*PCCA^KI/KI^* neonates received dual-gene AAV via facial-vein injection at postnatal day 1 (P1), after genotyping at P0, and plasma was sampled biweekly beginning at 4 weeks of age (Fig. 5A). Neonatal treatment significantly reduced glycine, the C3/C2 ratio, propionylglycine, 3-HP, and 2-MeCit compared with untreated controls (Fig. 5B-E, G), with propionylglycine falling to levels comparable to WT and heterozygous mice (Fig. 5D). Lysine and alanine did not differ significantly among groups (Fig. 5F, H).

**Figure 5.**
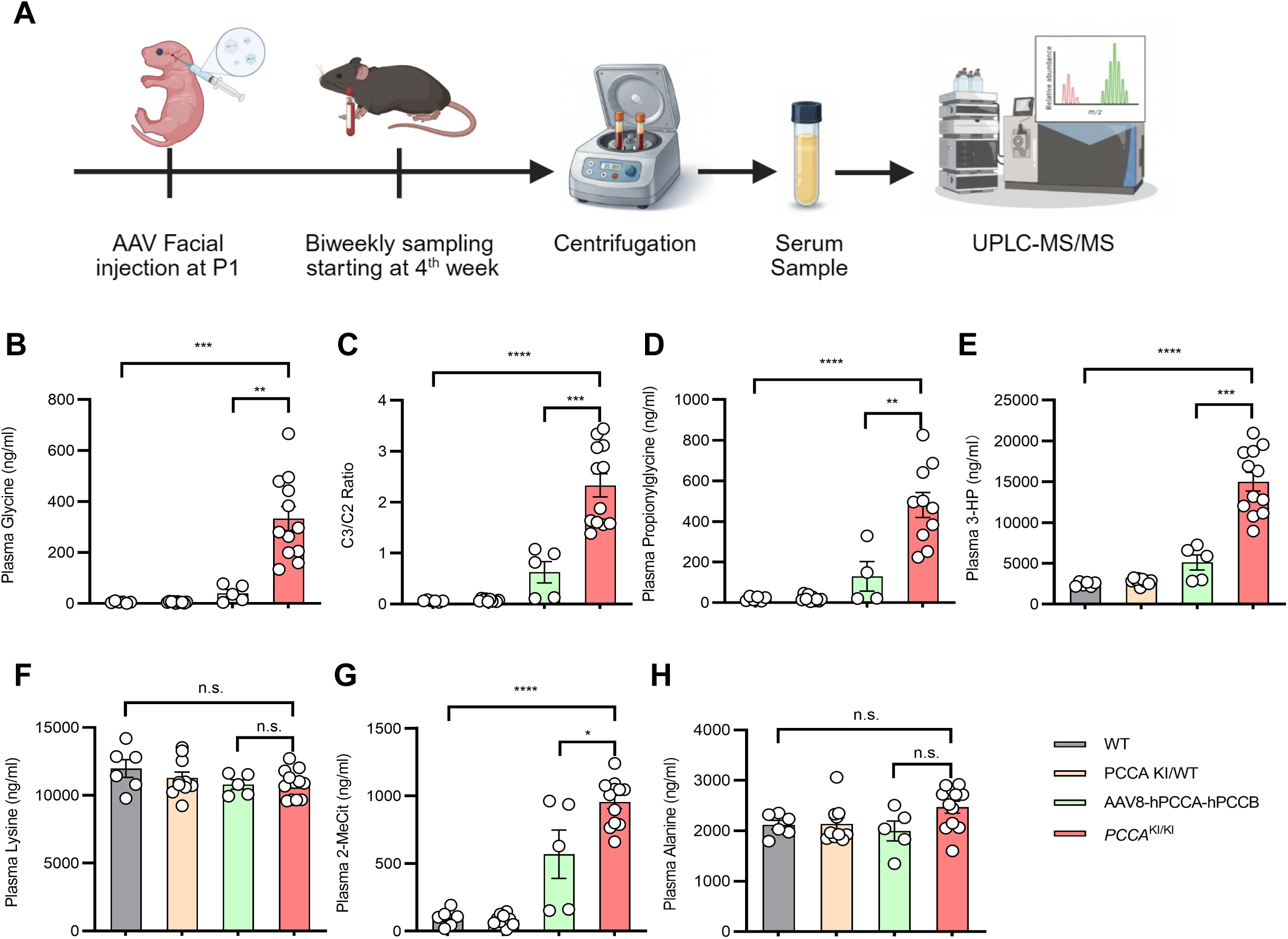
Neonatal administration of dual-gene AAV therapy achieved early and robust correction of circulating PA biomarkers. (A) Schematic of the neonatal treatment study design: *PCCA*^KI/KI^ pups were genotyped at postnatal day 0 (P0) and received a single facial-vein injection of AAV8-hPCCA-hPCCB at P1; plasma was sampled biweekly beginning at 4 weeks of age and analyzed by UPLC-MS/MS. (B-H) Plasma metabolite levels at 4 weeks post-injection for glycine (B), C3/C2 (propionylcarnitine/acetylcarnitine) ratio (C), propionylglycine (D), 3-hydroxypropionic acid (3-HP) (E), lysine (F), 2-methylcitric acid (2-MeCit) (G), and alanine (H). All metabolite concentrations are reported in ng/mL, except for the C3/C2 ratio, which is reported as a ratio. Data are presented as the mean ± SEM. \**p* < 0.05; \*\**p* < 0.01; \*\*\**p* < 0.001; \*\*\*\**p* < 0.0001; n.s., not significant (one-way ANOVA followed by Tukey’s posttest). Sample sizes are indicated in the corresponding panels.

To compare neonatal and adult treatment directly, we assessed both cohorts at 1-month post-injection: P1-treated mice were sampled at 4 weeks of age, and 4-week-treated mice at 8 weeks of age (the 8-week time point in Fig. 4A-G). Compared to the respective untreated *PCCA^KI/KI^* controls, P1 injection produced greater reductions in glycine (88.0% vs 79.8%), 2-MeCit (40.5% vs 3.7%), 3-HP (65.9% vs 56.3%), and the C3/C2 ratio (73.2% vs 69.4%), while propionylglycine correction was comparable between the two regimens (73.0% vs 76.9%). The advantage was most striking for 2-MeCit, where neonatal treatment achieved more than ten-fold greater reduction than adult dosing. These data support the case for early postnatal intervention, when lower baseline metabolite burden may favor a stronger therapeutic response.

## Discussion

Propionic acidemia remains a serious inborn metabolic disorder for which dietary restriction, pharmacological support, and liver transplantation fail to restore the missing enzymatic activity or prevent long-term disease progression[4,5,20,21]. Here, we show that a single intravenous injection of an AAV8 vector with a liver-specific promoter encoding both human *PCCA* and *PCCB* normalizes circulating PA biomarkers in a clinically relevant knock-in mouse model, with the effect persisting for at least three months four months and no detectable organ toxicity.

The rationale for delivering both subunits is straightforward: PCC is a heterododecameric complex in which *PCCB* is unstable without *PCCA* [1,8]. Replacing one subunit alone is unlikely to restore balanced stoichiometry or full activity across both PA subtypes. Our data bear this out. Single-gene AAV-h*PCCA* produced only partial correction and failed to reduce 2-MeCit or alanine, whereas co-delivery of both subunits achieved broad biochemical rescue (Fig. 3). This is consistent with the dual mRNA findings of superior efficacy with two-subunit delivery [11]. The key difference is durability: their mRNA approach required repeated dosing and did not fully normalize biomarkers even after 6 months[12,13], while a single AAV injection maintained metabolic correction over our entire follow-up period. For a chronic pediatric disease requiring lifelong management, this distinction matters.

We chose a humanized *PCCA*^R73W^ knock-in model rather than a conventional knockout[22,23]. The R73W substitution, modeling the human p.R77W variant[8], preserves residual PCC activity and produces a chronic, non-lethal phenotype that more faithfully tracks the metabolic course of patients with PA.. This allowed us to follow the treated mice for 16 weeks without confounding neonatal mortality. We also found that native human *PCCA*/*PCCB* sequences outperformed codon-optimized variants in vivo (Fig. S3A, B), a finding with practical implications for vector design that deserves further mechanistic study.

Notably, P1 facial-vein injection, modeling the earliest feasible window after newborn screening, achieved metabolic correction that was comparable to or better than that achieved by 4-week adult treatment (Fig. 5). This argues for early intervention, when hepatocyte proliferation may favor AAV transduction[24] and before irreversible organ damage has set in. PA patients depend on strict protein restriction and remain at risk of decompensation during illness, growth, or dietary lapses[5,25]; the robust neonatal correction observed here suggests that early gene therapy could substantially reduce this lifelong vulnerability.

An important question for future studies is whether the sustained metabolic correction observed here translates into the reversal of subcellular mitochondrial pathology. Preliminary TEM data from our *PCCA*^KI/KI^ model shows marked cristae disruption in untreated mice (Fig. 1C), and ongoing studies are examining whether dual-gene AAV therapy restores mitochondrial ultrastructure and respiratory function in treated animals. Resolving this question will be critical for establishing whether gene therapy can address the organelle-level damage that underlies PA pathophysiology[26,27].

Several limitations should be noted. First, the *PCCA*^KI/KI^ model expresses a mutant *PCCA* protein antigenically indistinguishable from the transgene-derived wild-type *PCCA*, precluding direct quantification of exogenous protein by western blot; we plan to use ddPCR for vector genome and human-specific transcript measurement. Second, PCC enzymatic activity was not measured directly because conventional radioisotope assays were unavailable; an LC-MS/MS-based assay is under development. Third, while our 16-week follow-up showed no waning of efficacy, longer observation is needed to assess AAV transgene persistence through hepatocyte turnover in a growing liver. Fourth, immune responses to the P2A peptide and human PCC subunits in a murine host were not comprehensively evaluated[28,29] and will require large-animal studies. Despite these gaps, the data establishes proof of concept for a single-dose, dual-gene AAV therapy that achieves sustained metabolic correction and an acceptable safety profile in a clinically relevant PA model, warranting further development for clinical application in both type I and type II propionic acidemia.

## Materials and Methods

### Animal models and in vivo interventions

A clinically relevant *PCCA*^R73W^ knock-in (*PCCA*^KI/WT^) PA model and a *PCCA*^KO/KO^ strain were generated using CRISPR/Cas9. Native human *PCCA* and *PCCB* sequences were packaged into AAV8 vectors driven by a liver-specific TBG promoter, yielding single-gene and dual-gene therapeutic constructs. Vectors were systemically administered to *PCCA*^KI/KI^ mice at neonatal (P1) or juvenile (4 weeks) stages. All mice were housed under specific pathogen-free (SPF) conditions in a mouse facility. Animals in the experimental groups were sex- and age-matched. The animals used in the study complied with all relevant ethical regulations regarding animal research. All experimental protocols conducted in this study involving animals were approved by the Institutional Animal Care and Use Committee of Zhejiang University.

### Biochemical, cellular, and histological assessments

Dual-gene expression was validated in vitro using lentivirus-transduced *PCCA*^KO/KO^ mouse embryonic fibroblasts. Plasma PA biomarkers (e.g., C3/C2 ratio, 3-HP, and 2-MeCit) were quantified via UPLC-MS/MS. Safety profiles and organ histology were assessed via serum biochemical panels, hematoxylin and eosin staining, and transmission electron microscopy (TEM).

### Statistical analysis

Data are expressed as mean ± SEM. Statistical significance was determined using an unpaired two-tailed Student’s t-test or analysis of variance (ANOVA) with Tukey’s post hoc test.Comprehensive methodological details are provided in the Supplementary Materials.

## Data Availability Statement

All data relevant to the study are included in the article or uploaded as supplementary information. Further data are available from the corresponding authors upon request.

## Author Contributions

H.X., X.H., and Y.Z. conceived the study and designed the experimental framework. X.H. established the clinical cohort, managed patient samples and medical records, and provided clinical consultation. H.X. and Z.T. performed the in vitro experiments. H.X. conducted the in vivo studies, including animal handling and tissue collection. J.L. and H.Z. were responsible for the generation and purification of AAV vectors. A.M. refined the manuscript to meet academic English standards and interpreted the mitochondrial function data. S.Y. and D.C. designed and generated the *PCCA* KI mouse model, which was generously provided by D.L.. The original manuscript was drafted, reviewed, and edited by H.X., Z.T., and Y.Z., with critical revisions and input from all authors. L.A. and Y.Z. secured funding for the project. L.A., X.H., and Y.Z. provided overall supervision for the research.

## Acknowledgements

This work was supported by the National Natural Science Foundation of China (82371861, 82402020), Key R&D Program of Zhejiang (2024SSYS0020), Henan Province Key Research and Promotion Project (242102311023), the Zhejiang Provincial Leading Innovation and Entrepreneurship Team Introduction and Cultivation Program (2024R01024), and the Starting Fund from Zhejiang University. We thank Shuangwei Hu from Xbiome Co. Ltd for advice on sequence optimization. We thank the core facility of Liangzhu Laboratory for the technical support.

## Supplementary Figure Legends

**Figure S1. Generation and additional phenotyping of the humanized *PCCA*^R73W^ knock-in mouse model.**

(A) Alignment of the human *PCCA* c.229C>T (p.R77W; CGG>TGG) variant with the corresponding murine *Pcca* R73W (AGG>TGG) substitution introduced to model the clinically relevant mutation. (B) Schematic of CRISPR/Cas9-mediated genome editing used to generate the *PCCA*^R73W^ knock-in allele (*PCCA*^KI/WT^), yielding an Arg-to-Trp substitution at the encoded protein level. (C) Blood ammonia levels in WT and *PCCA* knock-in mice, including sex-stratified measurements for *PCCA*^KI/KI^ animals as indicated. (D) Transmission electron microscopy of cardiomyocytes demonstrating mitochondrial ultrastructural abnormalities in *PCCA*^KI/KI^ mice compared with WT, with representative regions highlighted (dashed outline). (E) Longitudinal quantification of plasma propionylglycine showing marked accumulation in *PCCA*^KI/KI^ mice relative to WT and heterozygous controls. (F-H) Targeted metabolite profiling demonstrating increased plasma amino acids in *PCCA*^KI/KI^ mice, including glycine (F), alanine (G), and lysine (H). Data points represent group averages ± SEM. \**p* < 0.05; \*\**p* < 0.01; \*\*\**p* < 0.001; \*\*\*\**p* < 0.0001 (one-way ANOVA followed by the Tukey’s posttest).

**Figure S2. Validation of MEF identity and confirmation of transgene mRNA expression following lentiviral transduction.**

Immunofluorescence staining for vimentin (green) and DAPI (blue) in MEFs derived 510 from WT, *PCCA*KO/KO, and *PCCAKI/KI* mice, confirming mesenchymal identity across all 511 genotypes. (B, C) Quantitative RT-PCR analysis of relative *PCCA* (B) and *PCCB* (C) mRNA expression in PCCAKO/KO MEFs transduced with Lenti-hPCCA/B compared with untransduced PCCAKO/KO controls, demonstrating robust transgene-derived mRNA expression for both subunits.

**Figure S3. Native human *PCCA*/*PCCB* sequences outperform codon-optimized variants, and dual-gene AAV therapy sustains dose-dependent metabolic correction at 1 and 2 months post-treatment.**

(A, B) Comparison of codon-optimized (opti-*PCCA*) versus native non-codon-optimized (unopti-*PCCA*) human *PCCA*/*PCCB* sequences delivered via AAV8 in *PCCA*^KI/KI^ mice, assessed by plasma C3/C2 ratio (A) and 3-hydroxypropionic acid (3-HP, ng/mL) (B). Native sequences produced significantly greater reductions in both biomarkers compared with codon-optimized sequences. WT, *PCCA*^KI/WT^, and untreated *PCCA*^KI/KI^ are shown as controls. (C, D) Dose-response analysis of dual-gene AAV therapy 1×10¹¹, 5×10¹¹, and 1×10¹² vg/mouse) at 1 month post-injection on plasma alanine (C, ng/mL) and 2-methylcitric acid (2-MeCit) (D, ng/mL). (E-J) Extended metabolite profiling at 2 months post-injection across the same three dose groups for plasma glycine (E), alanine (F), lysine (G), propionylglycine (H), C3/C2 ratio (I), and 2-MeCit (J). All metabolite concentrations are reported in ng/mL except C3/C2, which is reported as a ratio. Data are presented as mean ±SEM. \**p* < 0.05;\*\**p* < 0.01; \*\*\**p* < 0.001; \*\*\*\**p* < 0.0001; n.s., not significant (one-way ANOVA followed by Tukey’s posttest). Sample sizes are indicated in the corresponding panels.

**Figure S4. Extended safety monitoring confirms absence of organ toxicity at 8 and 12 weeks post-injection.**

(A-D) Serum safety biomarkers at 8 weeks post-injection (12 weeks of age): alanine aminotransferase (ALT, U/L) (A), aspartate aminotransferase (AST, U/L) (B), urea (mmol/L) (C), and creatine kinase (CK, U/L) (D), compared across WT, *PCCA*^KI/WT^, AAV8-h*PCCA*-h*PCCB* (dual-gene), AAV8-h*PCCA* (single-gene), and *PCCA*^KI/KI^ groups. (E-H) The same safety biomarkers at 12 weeks post-injection (16 weeks of age): ALT (E), AST (F), urea (G), and CK (H). No significant differences were observed among groups at either time point.Data are presented as mean ± SEM (one-way ANOVA followed by Tukey’s posttest). Sample sizes are indicated in the corresponding panels.

## References

[1] Wongkittichote P, Ah Mew N, Chapman KA. Propionyl-CoA carboxylase – A review. Molecular Genetics and Metabolism 2017;122:145–52. 10.1016/j.ymgme.2017.10.002.

[2] Ugarte M, Pérez-Cerdá C, Rodríguez-Pombo P, Desviat LR, Pérez B, Richard E, et al. Overview of mutations in the PCCA and PCCB genes causing propionic acidemia. Hum Mutat 1999;14:275–82. 10.1002/(SICI)1098-1004(199910)14:4%253C275::AID-HUMU1%253E3.0.CO;2-N.

[3] Huang CS, Sadre-Bazzaz K, Shen Y, Deng B, Zhou ZH, Tong L. Crystal structure of the alpha(6)beta(6) holoenzyme of propionyl-coenzyme A carboxylase. Nature 2010;466:1001–5. 10.1038/nature09302.

[4] Marchuk H, Wang Y, Ladd ZA, Chen X, Zhang G-F. Pathophysiological mechanisms of complications associated with propionic acidemia. Pharmacology & Therapeutics 2023;249:108501. 10.1016/j.pharmthera.2023.108501.

[5] Baumgartner MR, Hörster F, Dionisi-Vici C, Haliloglu G, Karall D, Chapman KA, et al. Proposed guidelines for the diagnosis and management of methylmalonic and propionic acidemia. Orphanet J Rare Dis 2014;9:130. 10.1186/s13023-014-0130-8.

[6] Almási T, Guey LT, Lukacs C, Csetneki K, Vokó Z, Zelei T. Systematic literature review and meta-analysis on the epidemiology of propionic acidemia. Orphanet J Rare Dis 2019;14:40. 10.1186/s13023-018-0987-z.

[7] Chapman KA, Gropman A, MacLeod E, Stagni K, Summar ML, Ueda K, et al. Acute management of propionic acidemia. Molecular Genetics and Metabolism 2012;105:16–25. 10.1016/j.ymgme.2011.09.026.

[8] Clavero S, Martınez MA, Pérez B, Pérez-Cerdá C, Ugarte M, Desviat LR. Functional characterization of PCCA mutations causing propionic acidemia. Biochimica et Biophysica Acta (BBA) - Molecular Basis of Disease 2002;1588:119–25. 10.1016/S0925-4439(02)00155-2.

[9] Ohura T, Kraus JP, Rosenberg LE. Unequal synthesis and differential degradation of propionyl CoA carboxylase subunits in cells from normal and propionic acidemia patients. Am J Hum Genet 1989;45:33–40.

[10] Lam Hon Wah AM, Lam KF, Tsui F, Robinson B, Saunders ME, Gravel RA. Assignment of the alpha and beta chains of human propionyl-CoA carboxylase to genetic complementation groups. Am J Hum Genet 1983;35:889–99.

[11] Jiang L, Park J-S, Yin L, Laureano R, Jacquinet E, Yang J, et al. Dual mRNA therapy restores metabolic function in long-term studies in mice with propionic acidemia. Nat Commun 2020;11:5339. 10.1038/s41467-020-19156-3.

[12] Baek R, Coughlan K, Jiang L, Liang M, Ci L, Singh H, et al. Characterizing the mechanism of action for mRNA therapeutics for the treatment of propionic acidemia, methylmalonic acidemia, and phenylketonuria. Nat Commun 2024;15:3804. 10.1038/s41467-024-47460-9.

[13] Koeberl D, Schulze A, Sondheimer N, Lipshutz GS, Geberhiwot T, Li L, et al. Publisher Correction: Interim analyses of a first-in-human phase 1/2 mRNA trial for propionic acidaemia. Nature 2024;630:E13–E13. 10.1038/s41586-024-07646-z.

[14] Wang D, Tai PWL, Gao G. Adeno-associated virus vector as a platform for gene therapy delivery. Nat Rev Drug Discov 2019;18:358–78. 10.1038/s41573-019-0012-9.

[15] Li C, Samulski RJ. Engineering adeno-associated virus vectors for gene therapy. Nat Rev Genet 2020;21:255–72. 10.1038/s41576-019-0205-4.

[16] Pérez-Cerdá C, Merinero B, Rodríguez-Pombo P, Pérez B, Desviat LR, Muro S, et al. Potential relationship between genotype and clinical outcome in propionic acidaemia patients. Eur J Hum Genet 2000;8:187–94. 10.1038/sj.ejhg.5200442.

[17] Chiu Y-H, Liu Y-N, Liao W-L, Chang Y-C, Lin S-P, Hsu C-C, et al. Two Frequent Mutations Associated with the Classic Form of Propionic Acidemia in Taiwan. Biochem Genet 2014;52:415–29. 10.1007/s10528-014-9657-6.

[18] Yan Z, Yan H, Ou H. Human thyroxine binding globulin (TBG) promoter directs efficient and sustaining transgene expression in liver-specific pattern [J. Gene 2012;506:289–94.

[19] Guenzel AJ, Hillestad ML, Matern D, Barry MA. Effects of Adeno-Associated Virus Serotype and Tissue-Specific Expression on Circulating Biomarkers of Propionic Acidemia. Human Gene Therapy 2014;25:837–43. 10.1089/hum.2014.012.

[20] Zhou G-P, Jiang Y-Z, Wu S-S, Kong Y-Y, Sun L-Y, Zhu Z-J. Liver Transplantation for Propionic Acidemia: Evidence From a Systematic Review and Meta-analysis. Transplantation 2021;105:2272–82. 10.1097/TP.0000000000003501.

[21] Pillai NR, Stroup BM, Poliner A, Rossetti L, Rawls B, Shayota BJ, et al. Liver transplantation in propionic and methylmalonic acidemia: A single center study with literature review. Mol Genet Metab 2019;128:431–43. 10.1016/j.ymgme.2019.11.001.

[22] Miyazaki T, Ohura T, Kobayashi M, Shigematsu Y, Yamaguchi S, Suzuki Y, et al. Fatal propionic acidemia in mice lacking propionyl-CoA carboxylase and its rescue by postnatal, liver-specific supplementation via a transgene *. Journal of Biological Chemistry 2001;276:35995–9. 10.1074/jbc.M105467200.

[23] Guenzel AJ, Hofherr SE, Hillestad M, Barry M, Weaver E, Venezia S, et al. Generation of a hypomorphic model of propionic acidemia amenable to gene therapy testing. Mol Ther 2013;21:1316–23. 10.1038/mt.2013.68.

[24] Foust KD, Nurre E, Montgomery CL, Hernandez A, Chan CM, Kaspar BK. Intravascular AAV9 preferentially targets neonatal neurons and adult astrocytes. Nat Biotechnol 2009;27:59–65. 10.1038/nbt.1515.

[25] Ehrenberg S, Walsh Vockley C, Heiman P, Ammous Z, Wenger O, Vockley J, et al. Natural history of propionic acidemia in the Amish population. Molecular Genetics and Metabolism Reports 2022;33:100936. 10.1016/j.ymgmr.2022.100936.

[26] Zong Y, Li H, Liao P, Chen L, Pan Y, Zheng Y, et al. Mitochondrial dysfunction: mechanisms and advances in therapy. Sig Transduct Target Ther 2024;9:124. 10.1038/s41392-024-01839-8.

[27] Subramanian C, Frank MW, Tangallapally R, Yun M, White SW, Lee RE, et al. Relief of COA sequestration and restoration of mitochondrial function in a mouse model of propionic acidemia. J of Inher Metab Disea 2023;46:28–42. 10.1002/jimd.12570.

[28] Costa Verdera H, Kuranda K, Mingozzi F. AAV Vector Immunogenicity in Humans: A Long Journey to Successful Gene Transfer. Mol Ther 2020;28:723–46. 10.1016/j.ymthe.2019.12.010.

[29] Ertl HCJ. Immunogenicity and toxicity of AAV gene therapy. Front Immunol 2022;13:975803. 10.3389/fimmu.2022.975803.

