## Supplementary Materials and Methods_FigureS1-S4 for "AAV-Mediated Dual-Gene Therapy Restores Metabolic Function in Mice with Propionic Acidemia"

##### Generation and Genotyping of *PCCA*$^{KI/KI}$ and *PCCA*$^{KO/KO}$ Mouse Models

The mutant alleles for both *Pcca* knock-in (KI) and knockout (KO) models were generated on a C57BL/6J genetic background utilizing CRISPR/Cas9-mediated genome editing. For the KI allele, a precise point mutation corresponding to the human *PCCA* p.R77W (c.229C>T) clinical variant was introduced into exon 3 of the murine *Pcca* locus, culminating in an Arg-to-Trp substitution (R73W, AGG>TGG). Conversely, the KO allele was established by targeting and deleting exons 2 through 8 of the *Pcca* gene to generate a definitive null allele. Heterozygous founders were backcrossed with wild-type C57BL/6J mice to establish stable germline-transmitted colonies. Subsequently, heterozygous mice (*PCCA*$^{KI/WT}$ and *PCCA*$^{KO/WT}$) were intercrossed to generate the homozygous (*PCCA*$^{KI/KI}$ and *PCCA*$^{KO/KO}$) mice utilized in the experiments.

Genomic DNA was isolated from postnatal toe or tail biopsies using standard Proteinase K digestion and isopropanol precipitation. Genotyping was performed utilizing EazyTaq DNA Polymerase (Vazyme). For the KI locus, the target region was amplified (Forward: 5'-TAGTTGCCTTCTGGTTGTAAA-3'; Reverse: 5'-CTGTGTTTGTGATAACAGTGG-3') to yield a 533-bp product , with the specific point mutation subsequently verified via Sanger sequencing. For the KO locus, a multiplex PCR strategy with target-specific primers was utilized to differentiate the mutant (321 bp) and wild-type (483 bp) alleles.

##### Vector Construction and Viral Production

For in vitro validations, a dual-gene lentiviral expression vector was strategically engineered. Target sequences were seamlessly assembled into the vector backbone, and lentiviral particles were subsequently generated by co-transfecting HEK-293T cells with the transfer plasmid, packaging plasmid (psPAX2), and envelope plasmid (pMD2.G) using linear polyethylenimine. Viral supernatants were harvested at 48 and 72 hours post-transfection and filtered through a 0.45 μm membrane.

For in vivo therapeutic interventions, non-codon-optimized (native) human PCCA (NCBI Reference Sequence: NM_000282.4; CCDS9496.2) and PCCB (NCBI Reference Sequence: NM_001178014.2; CCDS54643.1) sequences were employed based on superior preliminary in vivo efficacy. Recombinant AAV8 vectors, governed by a liver-specific thyroxine-binding globulin (TBG) promoter, were manufactured via triple-plasmid transfection in HEK-293T cells. Vectors were rigorously purified utilizing iodixanol density gradient ultracentrifugation and titered via droplet digital PCR (ddPCR).

##### Primary Cell Isolation and In Vitro Assays

Mouse embryonic fibroblasts (MEFs) were successfully isolated from E14.5 embryos following enzymatic digestion with 0.25% Trypsin-EDTA. MEFs were maintained in High Glucose DMEM supplemented with 10% fetal bovine serum and 1% Penicillin-Streptomycin at 37°C within a humidified 5% CO2 environment.

For immunofluorescence evaluations, cultured cells were fixed in 4% paraformaldehyde and permeabilized/blocked using 5% normal donkey serum. Specimens were incubated overnight at 4°C with the following primary antibodies: mouse anti-PCCA (1:500; Santa Cruz), rabbit anti-PCCB (1:500; Proteintech), and rabbit anti-TOM20 (1:500; Proteintech). Following stringent washing, fluorophore-conjugated secondary antibodies (Goat anti-Mouse IgG 594 nm, Donkey anti-Rabbit IgG 546 nm, or Goat anti-Rabbit IgG Cy5 647 nm) were applied to visualize protein localization.

**Primary Hepatocyte Isolation and Mitochondrial Respiration Analysis**

At 12 weeks following AAV8 administration, primary hepatocytes were harvested utilizing an optimized two-step collagenase perfusion methodology. Mitochondrial respiratory dynamics were comprehensively profiled utilizing the Seahorse XFe96 Extracellular Flux Analyzer (Agilent Technologies). Hepatocytes were seeded into XFe96 microplates, permitting the dynamic quantification of the Oxygen Consumption Rate (OCR). Basal respiration, ATP-linked respiration, proton leak, and maximal respiratory capacities were calculated following sequential automated injections of oligomycin, FCCP, and a rotenone/antimycin A cocktail. Data visualization and statistical analyses were subsequently conducted utilizing GraphPad Prism.

**Neonatal Vector Administration and Dietary Protein Challenge**

To model the metabolic burden associated with propionic acidemia, heterozygous dams were acclimated to a 30% high-protein diet (HPD) prior to mating and maintained on this rigorous dietary regimen throughout gestation and lactation. Following parturition (P0), homozygous *PCCA*$^{KI/KI}$ neonates received a single systemic administration of the AAV8 vector at postnatal day 1 (P1). To facilitate early dietary adaptation, the HPD was introduced directly into the breeding enclosures at P12. All experimental pups were officially weaned at P21 and maintained continuously on the HPD. Longitudinal peripheral blood sampling for metabolite quantification commenced at P28.

**Targeted Metabolomic Profiling via UPLC-MS/MS**

Peripheral blood was collected at designated intervals, and plasma was rapidly isolated via centrifugation. The absolute concentrations of canonical PA biomarkers—specifically 3-hydroxypropionic acid (3-HP), 2-methylcitric acid (2-MeCit), propionylglycine, and a panel of relevant amino acids—were quantified utilizing an Ultra-Performance Liquid Chromatography system coupled with tandem mass spectrometry (UPLC-MS/MS). Target analytes were extracted via robust protein precipitation utilizing HPLC-grade methanol and acetonitrile, evaporated to complete dryness, and precisely reconstituted in the appropriate mobile phase prior to MS injection.

**Histopathological and Ultrastructural Analyses**

For histopathological safety assessments, target organs (heart, liver, lung, kidney) were excised, immediately fixed in 4% paraformaldehyde, embedded in paraffin, sectioned, and stained with hematoxylin and eosin (H&E). For transmission electron microscopy (TEM), mice were deeply anesthetized, and heart and liver tissues were rapidly harvested, minced into 1 $mm^3$ blocks, and submerged in 2.5% glutaraldehyde at 4°C overnight. Tissues underwent secondary fixation in osmium tetroxide, followed by graded dehydration and resin embedding. Ultrathin sections were contrasted utilizing uranyl acetate and lead citrate and examined under a 120 kV transmission electron microscope (Talos L120C, Thermo Fisher) to assess mitochondrial ultrastructural integrity.

**Nucleotide Sequences of Optimized Constructs**

The codon-optimized human sequences formulated for comparative *in vivo* evaluations in this study are detailed below:

**Codon-optimized human PCCA nucleotide sequence:**

ATGGCAGGATTCTGGGTAGGAACCGCTCCACTTGTGGCAGCAGGCCGGAG
GGGCCGATGGCCCCCCCAGCAGCTCATGTTGTCTGCAGCCTTGAGAACCCT
GAAACACGTGTTGTACTATTCCAGGCAGTGTCTGATGGTAAGTAGAAATCT
GGGATCCGTGGGGTATGATCCAAACGAGAAGACTTTCGATAAAATCTTGGT
TGCTAACCGAGGAGAGATCGCCTGTAGAGTCATCCGGACCTGCAAAAAGA
TGGGAATCAAAACAGTGGCAATTCATAGCGACGTAGATGCCAGTAGTGTCC
ACGTGAAAATGGCTGACGAGGCTGTTTGTGTTGGACCCGCTCCCACTAGTA
AATCCTACCTGAATATGGACGCTATCATGGAAGCCATCAAGAAAACTAGGG
CACAGGCTGTGCATCCCGGCTATGGCTTTCTCTCTGAGAACAAAGAGTTTG
CACGCTGCCTTGCTGCCGAGGATGTTGTCTTTATCGGGCCCGATACCCATGC
CATCCAGGCAATGGGCGATAAGATTGAGTCCAAACTGCTGGCAAAAAAGG
CAGAAGTTAACACGATTCCTGGCTTTGACGGCGTTGTGAAGGACGCTGAG
GAAGCCGTCCGAATTGCCAGGGAGATCGGGTATCCCGTAATGATAAAAGCC
AGCGCTGGTGGCGGTGGAAAAGGTATGAGAATCGCTTGGGACGATGAAGA
GACACGAGATGGATTTAGACTGTCCAGCCAGGAGGCTGCTTCCTCATTCGG
CGACGATCGACTGCTGATCGAAAAGTTTATCGACAATCCACGCCATATCGA
GATCCAGGTGCTGGGCGACAAACATGGTAATGCACTTTGGCTCAACGAGC
GGGAATGTTCCATCCAACGGAGGAATCAGAAGGTGGTGGAAGAGGCTCCT
TCCATCTTTCTGGATGCCGAAACTCGCAGAGCAATGGGAGAACAGGCAGT

GGCTCTGGCAAGAGCCGTCAAGTATTCATCCGCTGGCACCGTGGAATTTCT
GGTGGATTCTAAGAAAAATTTTTACTTCCTGGAGATGAATACTCGCCTGCAG
GTGGAGCATCCTGTTACCGAGTGCATTACAGGTTTGGATCTGGTGCAAGAG
ATGATCCGAGTCGCAAAAGGATACCCCCTGAGGCACAAGCAGGCCGACAT
CAGGATCAACGGTTGGGCCGTTGAGTGTCGAGTGTATGCCGAGGACCCCTA
CAAGTCCTTTGGACTTCCCAGTATTGGTCGACTCTCTCAGTACCAGGAGCC
TTTGCACCTTCCAGGTGTGCGAGTGGATAGTGGGATTCAGCCTGGTTCCGA
CATCTCTATCTATTACGACCCAATGATAAGTAAGCTCATCACCTATGGCTCCG
ATAGAACCGAAGCTCTCAAAAGGATGGCTGACGCTCTGGACAATTATGTCA
TTCGGGGCGTGACACACAATATAGCCCTGTTGAGAGAGGTGATTATTAATTC
AAGGTTTGTCAAAGGGGATATAAGCACAAAGTTCCTTAGCGATGTCTATCC
CGACGGATTCAAGGGTCACATGCTGACAAAGTCCGAAAAGAATCAGCTTC
TGGCCATCGCATCCAGCCTCTTCGTGGCTTTCCAGTTGCGAGCCCAACACT
TTCAGGAGAATAGTCGGATGCCTGTGATAAAACCTGACATCGCCAACTGGG
AACTGAGCGTTAAACTCCATGACAAAGTGCACACCGTAGTAGCCTCTAATA
ATGGGAGCGTATTCAGTGTTGAAGTGGATGGGTCAAAGCTGAATGTTACCA
GCACGTGGAATCTGGCCTCCCCTCTGCTTTCCGTGAGCGTCGATGGCACAC
AACGGACTGTCCAGTGCCTGTCACGGGAGGCCGGTGGTAACATGAGCATT
CAGTTCCTGGGAACCGTGTATAAGGTCAATATACTGACCCGGCTGGCTGCA
GAGCTCAACAAGTTCATGCTGGAGAAGGTAACAGAAGATACCAGTAGTGT
CTTGAGGAGCCCCATGCCTGGCGTTGTGGTGGCCGTTAGCGTCAAACCTGG
CGATGCAGTGGCCGAAGGTCAGGAGATTTGTGTGATCGAGGCTATGAAGAT
GCAGAACTCTATGACCGCCGGTAAGACCGGCACCGTTAAGTCTGTGCATTG
CCAAGCAGGCGATACAGTGGGCGAGGGGGATCTCCTGGTCGAGCTTGAA

**Codon-optimized human PCCB nucleotide sequence:**

ATGGCTGCAGCTTTGAGGGTTGCCGCTGTGGGTGCAAGACTCAGTGTATT
GGCCTCCGGGCTGCGAGCCGCCGTAAGATCTCTGTGTTCACAGGCCACAT
CCGTAAACGAACGGATTGAAAACAAGAGACGCACCGCACTCTTGGGCGG
TGGCCAGAGACGAATTGACGCACAGCACAAAAGGGGAAAGCTGACAGCA
CGGGAGCGGATCTCACTGCTTCTGGACCCAGGATCTTTCGTGGAGTCAGA
TATGTTTGTCGAGCATCGATGCGCCGACTTTGGCATGGCTGCTGATAAGA
ACAAGTTCCCCGGTGATTCTGTGGTCACCGGAAGGGGCCGCATAAACGGG
AGATTGGTATATGTGTTCAGTCAACAGATCATCGGGTGGGCACAGTGGCT
CCCTCTTGTCATTTCCGCACTTTGGGAAGCCGAGGACTTCACCGTGTTCGG
CGGCTCTTTGAGTGGCGCCCACGCCCAGAAGATTTGCAAGATTATGGACC
AGGCTATAACAGTGGGAGCACCAGTGATTGGCCTGAACGATTCCGGAGGC
GCCAGAATTCAAGAGGGAGTTGAGTCACTCGCAGGATATGCCGATATCTT
CCTGCGCAATGTGACCGCATCTGGAGTGATTCCTCAGATTAGCCTGATAAT
GGGCCCTTGTGCCGGTGGTGCCGTCTACTCTCCAGCTTTGACTGATTTCAC
CTTTATGGTGAAAGACACTAGCTACCTCTTCATCACCGGCCCTGATGTCGT
CAAAAGCGTTACAATGAGGACGTCACTCAGGAGGAACTGGGGGGTGCA

AAAACCCACACTACCATGAGCGGCGTGGCTCACCGGGCATTTGAAAATGA
CGTTGACGCCTTGTGCAACCTCCGCGATTTCTTCAATTACCTTCCCCTGTCT
TCCCAAGACCCTGCACCTGTGAGAGAATGCCACGACCCCTCAGACCGACT
GGTGCCTGAACTTGACACCATAGTGCCTCTGGAGAGCACCAAGGCCTACA
ACATGGTGGACATCATCCATAGCGTGGTTGACGAACGCGAGTTTTTCGAG
ATTATGCCCAATTACGCCAAGAATATCATCGTTGGGTTCGCCCGCATGAAT
GGACGAACCGTCGGTATAGTGGGGAATCAGCCAAAGGTGGCCTCCGGGT
GTCTGGACATCAATTCATCCGTTAAGGGCGCCAGGTTCGTCCGGTTCTGCG
ATGCATTCAACATCCCCCTCATCACATTCGTGGATGTGCCAGGCTTCCTTC
CTGGCACTGCCCAGGAATATGGAGGTATAATTCGCCACGGCGCCAAGCTT
CTTTATGCATTTGCCGAGGCCACTGTCCCTAAGGTGACCGTCATTACACGA
AAAGCCTACGGCGGAGCATATGACGTGATGAGCTCTAAACATCTTTGTGG
AGATACAAACTATGCTTGGCCAACTGCTGAAATTGCAGTGATGGGCGCTA
AAGGTGCTGTGGAAATCATCTTCAAAGGTCACGAAAACGTAGAGGCAGC
ACAGGCAGAGTACATAGAAAAGTTCGCTAACCCTTTTCCCGCCGCCGTAA
GAGGTTTTGTTGACGACATAATCCAGCCTAGCAGTACTAGAGCCCGCATC
TGTTGTGATCTGGACGTCCTGGCAAGCAAAAAAGTACAGAGACCTTGGCG
AAAGCATGCTAACATCCCTTTGtaa

•

### Supplementary Figure 1

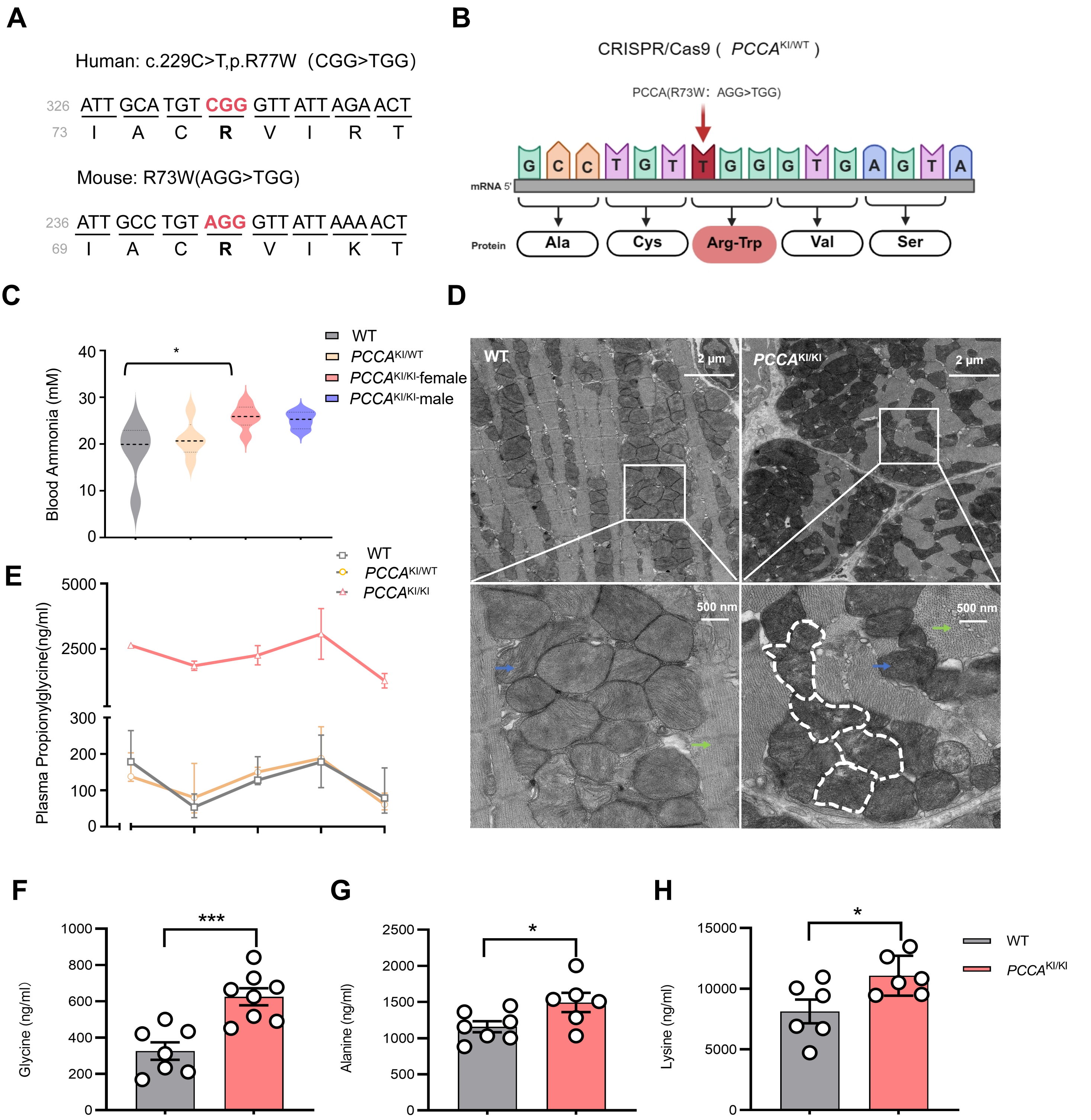

### Supplementary Figure 2

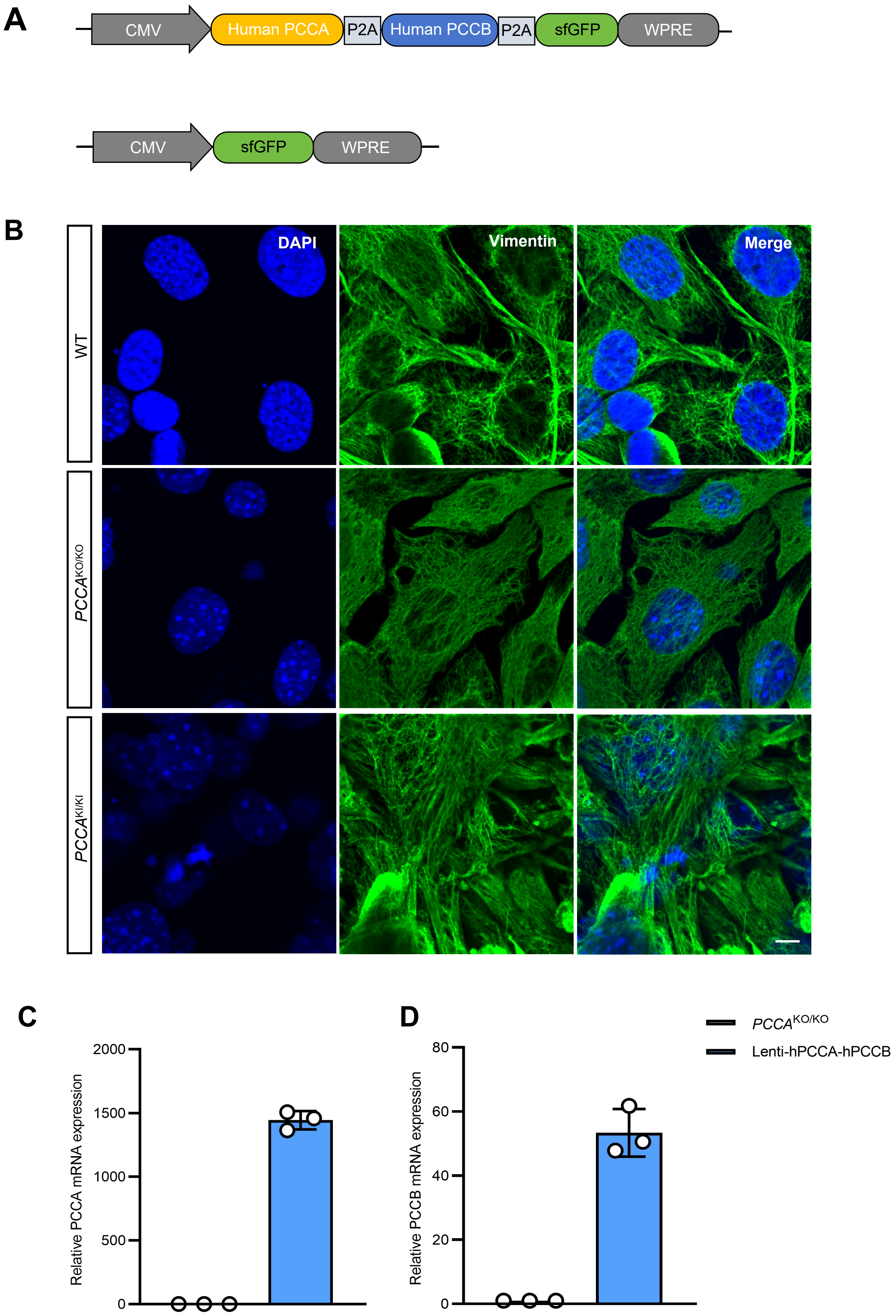

### Supplementary Figure 3

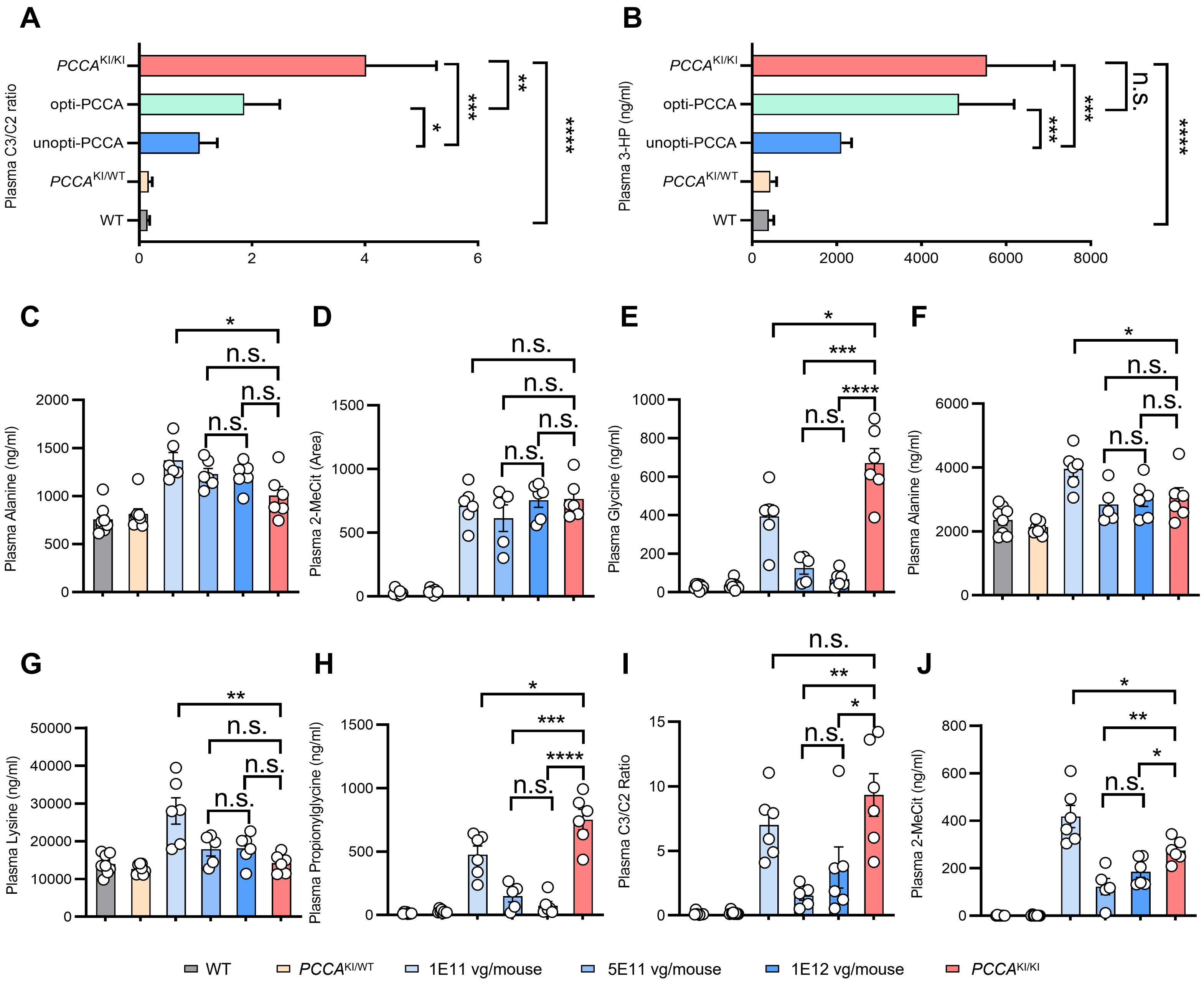

### Supplementary Figure 4

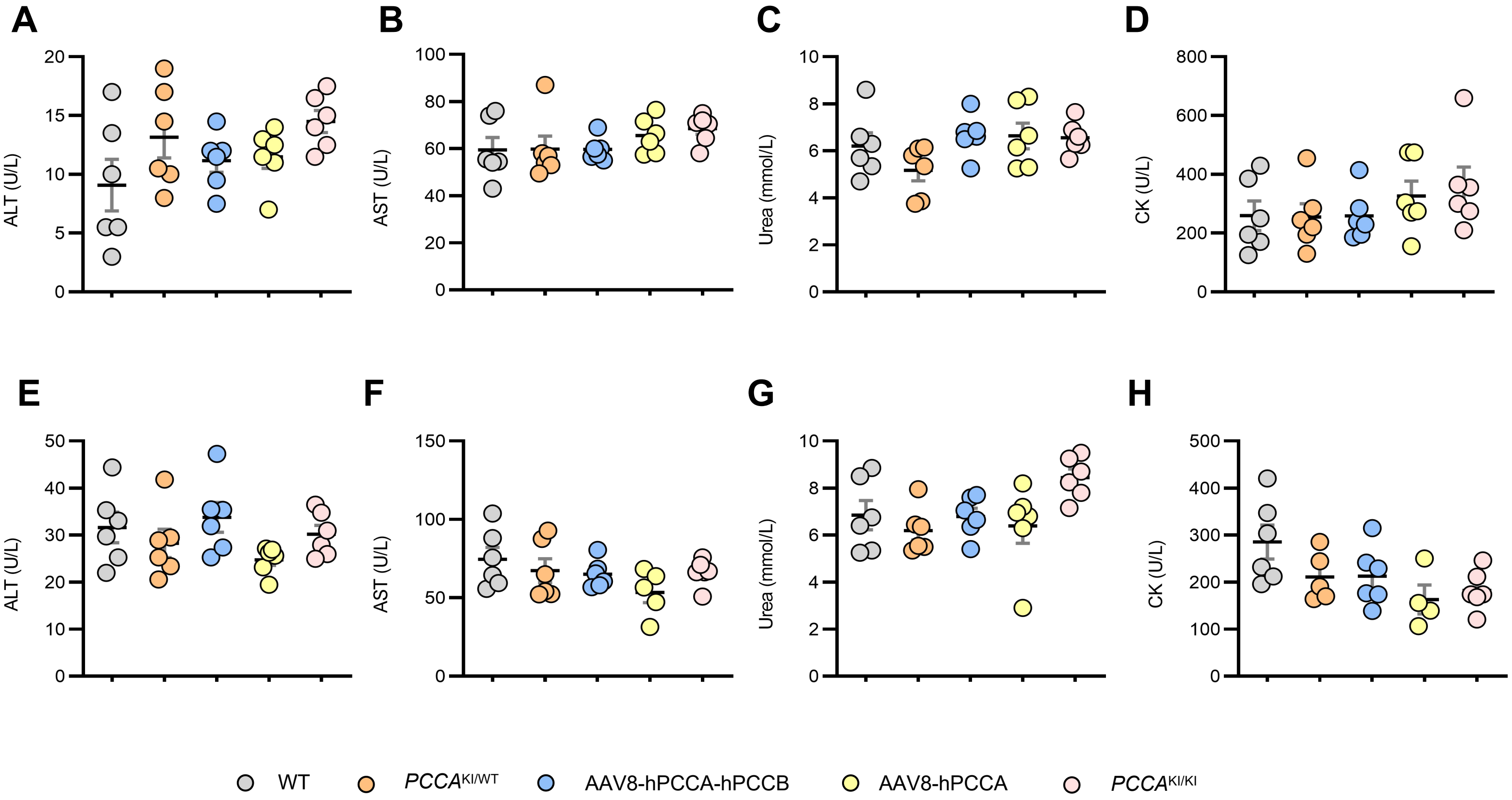
